# Exploring Resistance to ETS Targeting Agents in Diffuse Large B-Cell Lymphoma

**DOI:** 10.1101/2025.09.18.675797

**Authors:** Filippo Spriano, Luciano Cascione, Chiara Tarantelli, Giulio Sartori, Alberto J. Arribas, Adriana Velasova, Sara Napoli, Ondrej Havranek, Jeffrey A. Toretsky, Francesco Bertoni

**Affiliations:** Institute of Oncology Research, Faculty of Biomedical Sciences, USI, Bellinzona, Switzerland; SIB Swiss Institute of Bioinformatics, Lausanne, Switzerland; Biocev, First Faculty of Medicine, Charles University, Prague, Czech Republic; First Department of Medicine - Hematology, First Faculty of Medicine, Charles University and General University Hospital, Prague, Czech Republic; Departments of Oncology and Pediatrics, Lombardi Comprehensive Cancer Center, Georgetown University, Washington, DC, USA; Oncology Institute of Southern Switzerland, Ente Ospedaliero Cantonale, Bellinzona, Switzerland

**Author notes:** **Corresponding author:** Prof. Francesco Bertoni, Institute of Oncology Research, Faculty of Biomedical Sciences, USI, Bellinzona, via Francesco Chiesa 5, 6500 Bellinzona, Switzerland.

## Abstract

Diffuse large B-cell lymphoma (DLBCL) remains a challenging disease with limited therapeutic options beyond standard immunochemotherapy. ETS transcription factors, including SPIB and SPI1, are implicated in lymphoma pathogenesis and can be targeted by the small molecule TK216, which disrupts ETS–DHX9 interactions. To explore mechanisms of resistance, we generated stable TK216–resistant clones from the ABC-DLBCL line U2932. Resistant clones displayed a 4–5-fold increase in IC_50_ values and lost G2–M arrest upon treatment. Transcriptomic and mutational analyses revealed three resistance patterns: (i) MDR1/ABCB1 overexpression, leading to multidrug efflux; (ii) Cluster A, enriched for proliferation, Wnt, and transcriptional programs, with mutations in ESR2, USP24, and SFSWAP; and (iii) Cluster B, characterized by actin/microtubule remodeling, altered metabolism, and mutations in SRSF11 and PATJ. Pharmacologic screening identified increased sensitivity of resistant cells to BCL2, MCL1, and XPO1 inhibitors, while showing reduced sensitivity to aurora kinase and microtubule-targeting agents. Venetoclax and selinexor retained activity in resistant models, supporting their potential for rational combinations with TK216. These findings demonstrate that multiple, heterogeneous mechanisms drive resistance to ETS inhibition in DLBCL and highlight therapeutic strategies to overcome it.

## INTRODUCTION

Diffuse large B-cell lymphoma (DLBCL) is the most common lymphoma subtype, accounting for approximately 30-35% of all cases ^1^. Despite the essential improvements achieved in managing DLBCL patients ^2-4^, there is a need for therapies with novel mechanisms of action. Targeting ETS (E26 transformation-specific) factors is supported by the notion that this family of transcription factors plays a crucial role in regulating cell growth, differentiation, and survival across various cell types and tissues ^5^. In the context of DLBCL, dysregulation of ETS1, SPIB, and FLI1, members of the ETS transcription factor family, has been associated with DLBCL pathogenesis ^6-12^. Based on YK-4-279, TK216 is a first-generation clinical-grade small molecule that targets the EWS-FLI1 fusion protein ^13,14^. TK216 showed some clinical activity in Ewing sarcoma patients as a single agent or combined with vincristine; however, optimization is required, including an oral formulation ^15^. We have previously reported potent *in vitro* and *in vivo* anti-tumor activity of TK216 in DLBCL models ^13^. In DLBCL cells, TK216 exerts its activity, disrupting the interaction between the ETS factors SPIB or SPI1 and the RNA helicase DHX9 in activated B-cell-like (ABC) or germinal center B cell-like (GCB)-DLBCL, respectively ^13^. To better understand the mechanism of action of TK216 in DLBCL cells and identify ways to improve its efficacy, we have developed models of primary resistance to the compound.

## MATERIALS AND METHODS

### Cell lines and drug treatments

All cell lines used in this project were cultured under standard conditions (37 °C and 5% CO_2_ in a humidified atmosphere). Cell lines were obtained and supplemented with the appropriate medium, plus 10% Fetal Bovine Serum (FBS) (FBS-11A, Capricorn Scientific, Germany) and 1% penicillin/streptomycin (15070063, Thermo Fisher Scientific, Switzerland). All cell lines were periodically tested to confirm Mycoplasma negativity using the MycoAlert Mycoplasma Detection Kit (Lonza, Visp, Switzerland). In addition, all cell lines were validated for their identity by short tandem repeat (STR) DNA fingerprinting at IDEXX BioResearch (Ludwigsburg, Germany) or with the Promega GenePrint 10 System kit (B9510).

*In vitro* anti-proliferative activity was determined by treating cells for 72 hours. After 3 days, the MTT solution was added to the plates. After 4 hours, the reaction was stopped with SDS (25% solution), and absorbance was detected with Cytation 3 (Agilent, United States). IC50 calculations were done using an in-house R script. Live-cell-imaging treatments were done using an Incucyte machine (Sartorius); 30,000 cells per well were seeded in a 96-well plate and treated with TK216 (350nM), Tariquidar (5 μM), or Zosuquidar (5 μM) in single or combination and followed for approximately 4 days. TK216, YK-4-279, and tariquidar were purchased from MedChemExpress, zosuriquidar from LubioBioscience, and all the other compounds from Selleckchem.

### Resistance induction

To induce resistance to TK216, an ABC-DLBCL cell line, U2932, was seeded in a 24-well plate at a concentration of 50,000 cells / mL for each well and treated with an IC90 concentration (500nM) of TK216. In parallel, as a control, three flasks of U2932 were treated with the same volume of DMSO. After cells recovered from the treatment, they were treated periodically for almost one year, until the development of clones with stable TK216 resistance. We assessed the cell lines’ identity to determine whether contamination with other cells could have occurred during the continuous passages. Resistant and parental cell lines were tested by short tandem repeat DNA fingerprinting using the Promega GenePrint 10 System kit (B9510), showing that all clones derived from the parental U2932 cell lines.

### Combination studies

All combinations were performed by treating cells for 72 hours with increasing concentrations of both compounds in a 7 x 7 matrix in three technical replicates, followed by MTT assay and SDS cell lysis. The combination index (CI) was calculated using the Chou-Talalay method ^16^ and the HSA, ZIP, Loewe, and Bliss methods ^17^’

### Cell cycle

After the desired time for treatment, cells were harvested and washed twice in PBS. The cell pellet was collected, resuspended in 1ml PBS, and fixed with 4 mL of 100% ethanol molecular grade (CAS: 64-17-5, VWR, Belgium) and added drop by drop to the cells under constant mild agitation. Cells were then conserved for at least one night at -20 °C. Subsequently, cells were washed with PBS + 1% FBS and incubated with propidium iodide for at least 10 mins in the dark at 37 °C. Data acquisition was performed using the flow cytometry BD FACSCanto system (BD Bioscience), and data were analyzed with FlowJo software (Treestar).

### RNA Extraction

Total RNA extraction was done using TRIZOL reagent (Invitrogen) following the manufacturer’s protocol. During RNA isolation, samples were purified from potential genomic DNA contamination by DNase treatment using an RNase-free DNase Kit (Qiagen, Hilden, Germany). RNA concentration was determined using the Nanodrop spectrometer (NanoDrop Technologies) at 260 nm.

### Transcriptome profiling and data mining of RNA-Seq for identification of resistance-associated mutations

RNA-seq reads quality was assessed with FastQC (v0.11.5), and low-quality reads/bases and adaptor sequences were removed using Trimmomatic (v0.35). The trimmed, high-quality sequencing reads were aligned using STAR ^18^, a spliced read aligner allowing sequencing reads to span multiple exons. On average, we aligned 85% of the sequencing reads for each sample to the reference genome (HG38). The HTSeq-count software package ^19^ was then used for gene-level expression quantification. Differential expression analysis was performed on gene-level read count data using the ‘limma’ pipeline ^20^on gene-level read count data. We first subsetted the data to genes with counts-per-million values greater than one in 3 or more samples. The data were normalized per sample using the ‘TMM’ method from the edgeR package ^21^, and transformed to log2 counts-per-million using the edgeR function ‘cpm’. Linear model analyses, with empirical-Bayes moderated estimates of standard error, were then used to identify genes whose expression was most associated with the phenotype of interest, and an FDR-adjusted P-value of <0.05 was set as a threshold for statistical significance.

Reads in fastq format were pre-processed with GATK version 3.5 to remove Illumina adapter sequences (analysis type –T ClipReads, -XF illumina.adapters.fa) and Phred-scaled base qualities of 10 and below (-QT 10), similarly to what was described previously ^22^. After GATK processing, reads were mapped to HG38 using STAR (basic 2-pass method). SAMtools ^23^ flagstat was used to compute the number and percent of reads mapped to the genome. PCR/optical duplicates were marked by Picard (http://broadinstitute.github.io/picard/). Base quality recalibration and indel realignment were performed using GATK ^24^. Variant calling was performed using MuTect version 1.1.4 ^25^ for parental/resistant pairs. The variants (single-nucleotide mutations and indels) were annotated using Annovar ^26^. Variant filtering was performed using the following criteria: exonic, not synonymous variants, somatic variants, and damaging variants, excluding variants found in repetitive regions, variants found in areas with poor coverage, and reported SNPs.

### Gene set enrichment analysis (GSEA)

Log2-transformed and normalized expression profiles were used in Gene Set Enrichment Analysis (GSEA) ^27^ to functionally annotate differentially expressed genes. Standard settings were used for either regular GSEA or preranked GSEA (when using a ranked list organized after fold change), and signatures with nominal p-values <0.05 and FDR <0.05 were considered statistically significant. The ssGSEA data obtained with Gene Set Enrichment Analysis (GSEA) ^27^ were used as input for the Cytoscape software. Heatmaps were obtained with the R pheatmap package.

### Multidimensional scaling (MDS) plot and Heatmap

Analyses were performed using the R environment (R Studio console; RStudio, Boston, MA). Pearson’s clustering method was used for heatmaps.

## RESULTS

### The ABC-DLBCL cell line U2932 develops resistance to TK216 via multiple ways

To induce resistance to TK216, one ABC-DLBCL cell line, U2932, and one GCB-DLBCL cell line, DOHH2, were seeded at time 0 into 24 wells and treated with the drug at 500 nM and 1.5 μM, corresponding to the respective IC90 concentration, determined by 72h drug treatment. After almost two months, only cells from five wells out of 24 in U2932 were recovered after the first TK216 treatment. None of the 24 DOHH2 replicates developed stable resistance to TK216. The U2932 resistant cells (defined as clones 2, 3, 4, 5, 9) were then passed and treated again with TK216 (500nM) every three days for at least six months. The resistance was stable, as shown by the MTT assay performed in cells kept in a drug-free medium for at least two weeks, and the cells presented a 4-5 fold increase in their IC50 values versus their parental counterpart (Figure 1A). The average IC50 changes were from 150 nM to 1 µM. In addition, cells became resistant to YK-4-279 (Figure 1B), the ETS inhibitor from which TK216 is derived ^28^, but not resistant to a drug with a completely unrelated mode of action, such as vorinostat (Supplementary Figure 1).

**Figure 1.**
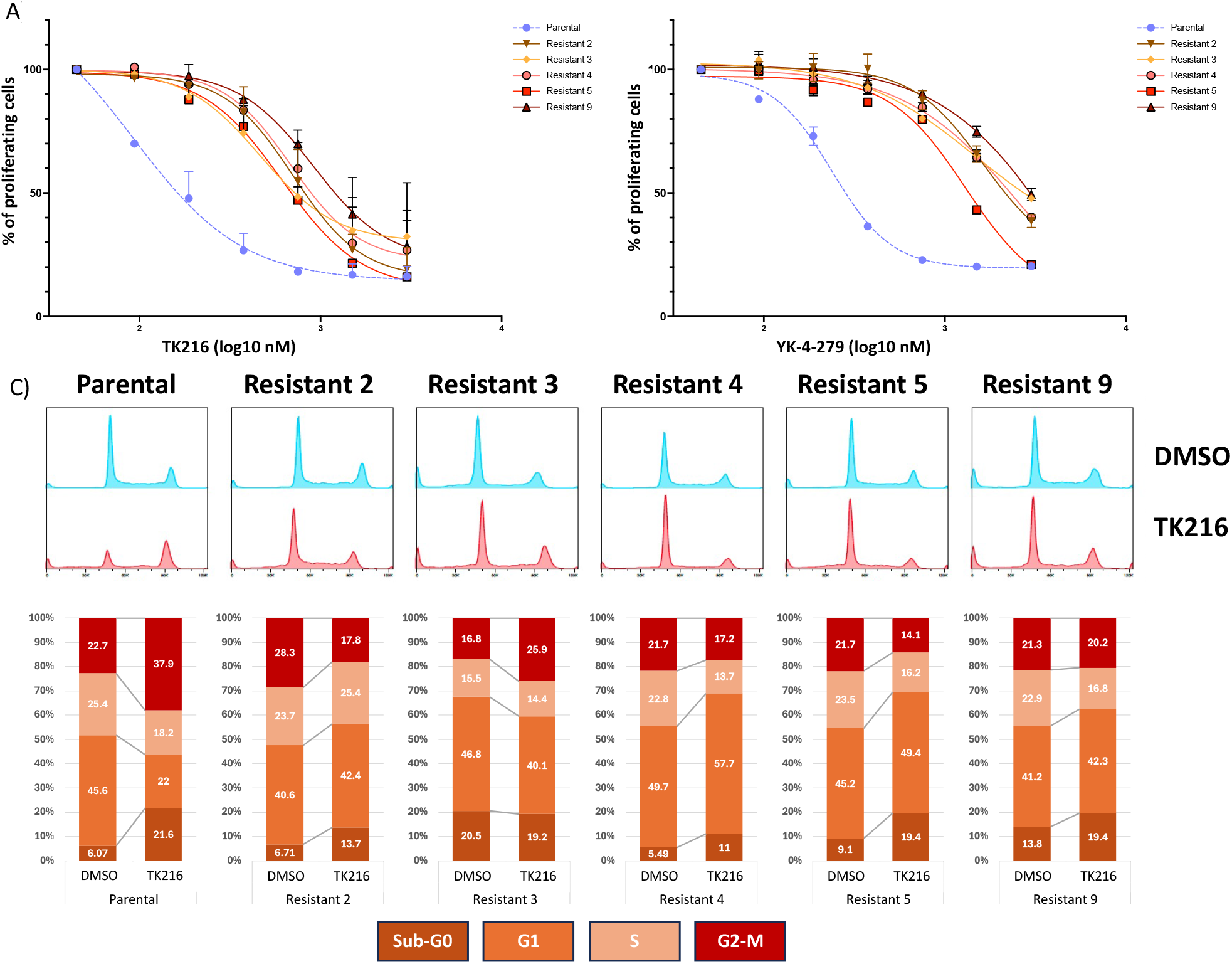
U2932 developed resistance to TK216. Resistant clones were treated with increasing doses of TK216 (A) or YK-4-279 (B) for 72 hours after 2 weeks of washout from the small molecule to validate the obtained resistance. The figure shows representative results from experiments performed in at least two replicates. C) Cell cycle performed in parental cell line compared to resistant clones after 24 hours of treatment with TK216, 500nM, or DMSO. The figure shows representative results from experiments performed in duplicate.

The acquisition of resistance was also demonstrated by performing a cell cycle analysis after 24h of TK216 treatment at 500 nM (IC90 of parental cell lines) (Figure 1C). The cell cycle analyses showed that resistant clones did not undergo G2-M cell cycle arrest but maintained a small subset, smaller than parental cells, of cell death, as represented by the small sub-G0 peak.

The five resistant clones and three biological replicates of the parental cell line underwent RNA-Seq to identify transcriptomic changes associated with decreased sensitivity to TK216 (Supplementary Table 1). The RNA extraction was performed in the absence of TK216 treatment for at least two weeks to observe stable differences. The MDS (multidimensional scaling) plot identified three main groups of cells: one with the three parental cell lines, one with Clone 2 and Clone 5, and one with Clone 3, Clone 4, and Clone 9 (Figure 2A). Focusing only on resistant cells, these were divided into three clusters: Clone 2 and 5 (Cluster A), Clone 3 and 4 (Cluster B), and Clone 9 (Figure 2B). We then performed a ssGSEA of parental and resistant cell lines, followed by an unsupervised clustering (Figure 2C). The clustering confirmed the MDS data, highlighting four clusters: parental, clones 2-5 (Cluster A), 3-4 (Cluster B), and clone 9. The detection of three clusters in resistant clones suggests three possible mechanisms that cells could have developed resistance to TK216.

**Figure 2.**
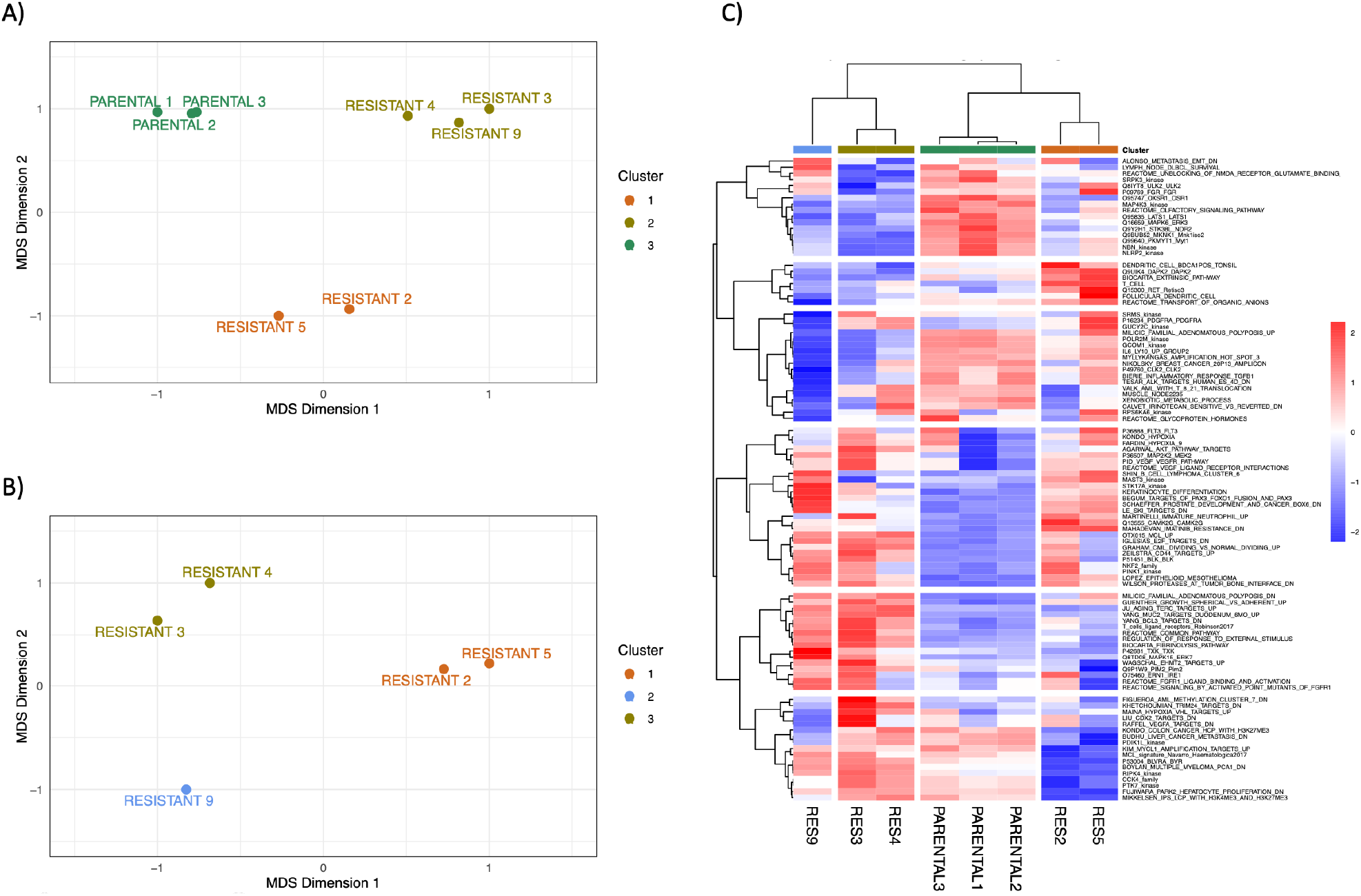
Multidimensional scaling (MDS) plot performed on RNA-Seq data on parental cell lines and resistant clones (A) and corresponding unsupervised clustering of ssGSEA (B). MDS plot based on RNA-seq data performed on A) parental and resistant clones and B) only in resistant clones. C) Unsupervised clustering of single-sample gene-sets enrichment data in parental and resistant cell lines. Pearson^’^s clustering method was used; the top 100 gene-sets for standard deviation are plotted.

### Resistance to TK216 can be due to multidrug resistance efflux pumps

To identify mechanisms underlying resistance, we first evaluated the expression levels of MDR genes that encode for drug efflux pump proteins. Based on RNA-Seq data, we identified MDR1 (ABCB1) as much more expressed in clone 9 than in parental cell lines and the remaining resistant clones (Figure 3A). MDR1 was also upregulated in clones 2, 3, and 4, but at a lower level than in clone 9. The data obtained with RNA-Seq were further validated by real-time PCR, highlighting a higher expression, normalized on parental levels, of MDR1, especially in clone 9. The upregulation was present in resistant clones under TK216 treatment selection and in clones with more than two weeks of drug washout, demonstrating that this is a stable mechanism that cells acquired during the time to survive TK216 exposure (Figure 3B) and not a transient mechanism upon TK216 exposure. The RNA data were further validated by immunoblot, showing an upregulation in MDR1 protein of 2 and 1.5-fold in clones 2 and 3, respectively, compared to parental cells, and a 6-fold increase in clone 9 (Figure 3C). Through live cell-imaging techniques, treatment of resistant cell lines (resistant 2, 3, and 9) with MDR1 inhibitors tariquidar and zosuquidar showed that clone resistant 9 showed a benefit from the combination treatments compared to single drug treatments (Figure 3D). This was also confirmed by the coefficient of drug interaction (CDI) <1 (Figure 3E). Finally, treatment with vincristine, a well-known substrate of different MDR proteins and especially of MDR1 ^29,30^, showed a significant increase in resistance in TK216- resistant clone 9, yet much smaller decreases in sensitivity in the remaining resistant clones (Figure 3F). These experiments further demonstrate how resistance clone 9 was driven by the efflux pump MDR1 overexpression.

**Figure 3.**
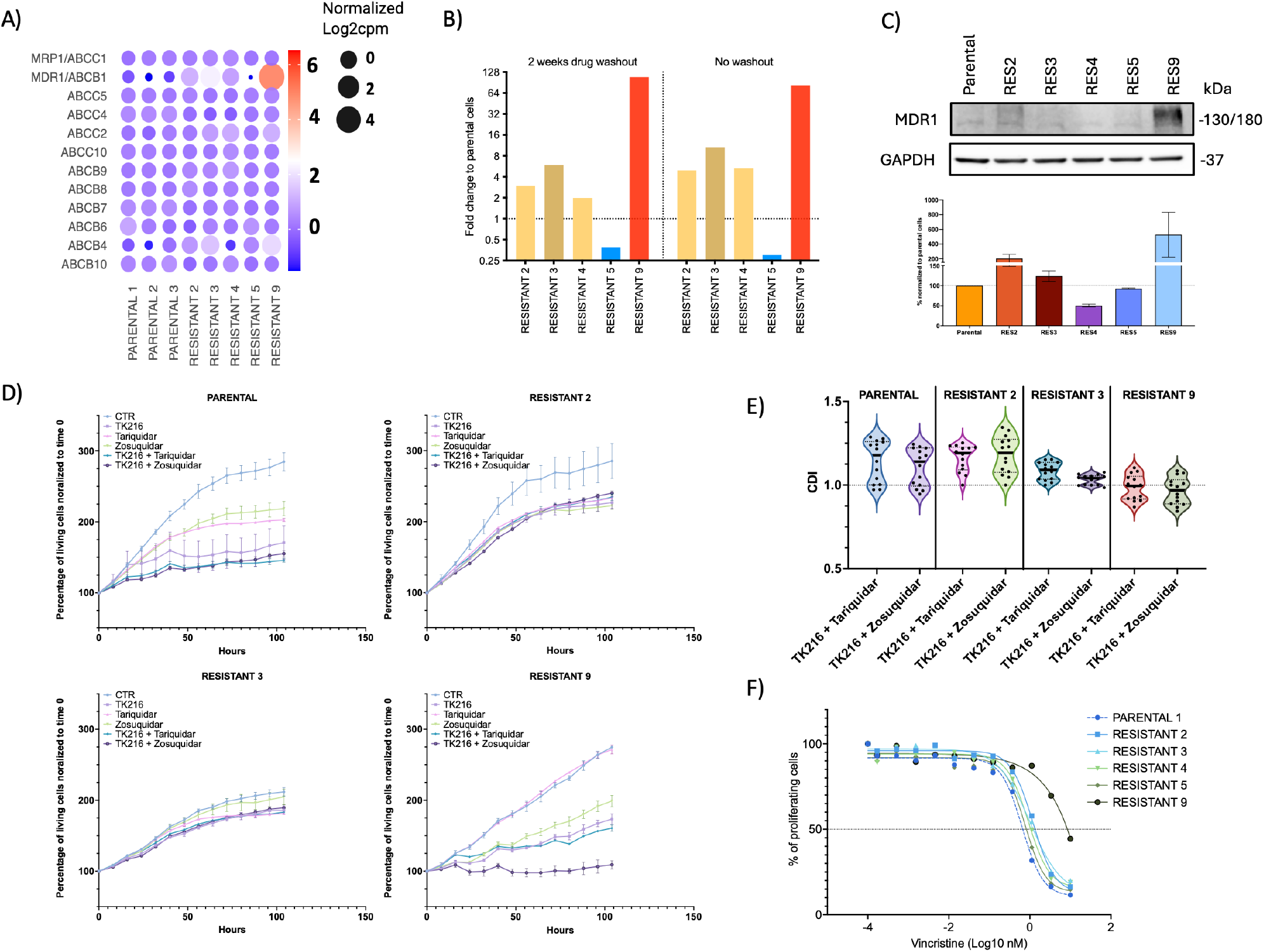
TK216 can be a substrate of multidrug resistance efflux pumps. A) Median normalized Log2 expression of genes coding for proteins involved in multi-drug resistance. B) MDR1 expression was obtained with real-time PCR and performed on resistant clones under TK216 or after 2 weeks of washout. Y-axis, Fold change between parental cell lines and resistant clones. C) Protein MDR1 expression with the respective quantification. Representative of two biological replicates. D) Growth curve of cells treated with TK216 or MDR1 inhibitors (tariquidar and zosuriquidar) in single or combination. E) Coefficient of drug interaction (CDI), CDI < 1 = synergism, CDI = 1 = additive, CDI > 1 = antagonism. F) Dose-response curve obtained after 72 hours of treatment with vincristine in parental and resistant clones. The figure shows representative results from experiments performed in duplicate.

### Characterization of Cluster A and Cluster B resistant clones

Clones 2 and 5 are analyzed together as Cluster A, and clones 3 and 4 are analyzed as Cluster B. In Cluster A, the top 50 genes whose expression was reduced compared to parental cells included genes involved in membrane organization and cell morphology (FMNL2, ACTN1, DBN1, and ADD2); Zinc finger proteins (ZNF57, ZNF470, ZBTB32, and ZNF792), and VASH1 (involved in microtubule dynamics). The top 50 genes, whose expression is increased in resistant cells, we identified genes involved in cell proliferation (SAMSN1, PITX2, MEOX2, and MECOM); genes involved in calcium binding or channel, such as SPOCK2, CNGB3, EFCAB12, TRPC4; genes involved in wnt pathway (TPB, LGR6, and CXXC4); genes involved in transcriptional regulation (TFEC, ZFP30, ZNF154, TWIST1, ID4, FOXP2, and MECOM) and genes involved in microtubule formation (MAP2, KIF13A, MAP7) (Supplementary Figure 2A).

In Cluster B, the top 50 genes whose expression was reduced compared to parental cells included genes involved in microtubule dynamics (FSD1, CAMSAP2, CENPE, MID1IP1, GPHN); Zinc finger proteins (ZNF354B, ZNF234, ZNF57), and the actin-related gene ACTN1. We identified actin-binding genes among the top 50 upregulated genes, such as SYNPO, DMTN, and FSCN2 (Supplementary Figure 2B).

Supplementary Figure 3 summarizes the gene sets enriched among the transcripts differentially expressed between resistant and parental cells. Transcripts upregulated in Cluster A were enriched in the hypoxia pathway, extracellular matrix-associated genes, interferon alpha pathway, secreted factors, genes upregulated after treatments with PI3K or BET inhibitors, GPCRs, genes upregulated after IRF4 knockdown, genes repressed by SPIB, and genes modulated after lenalidomide treatment. We also identified a downregulation in pathways such as DNA repair, cell cycle and mitosis, aurora kinase A activation, MYC targets, genes involved in the spliceosome and ribosome machinery, and genes involved in the B-cell receptor (BCR).

In Cluster B, upregulated transcripts, similarly to what was seen for Cluster A, were enriched in extracellular matrix-associated genes, genes modulated after lenalidomide treatment, genes upregulated after treatments with PI3K or BET inhibitors, and GPCRs. In contrast, unlike Cluster A, genes were downregulated after IRF4 or SPIB knockdown, and metabolism pathways were upregulated. Among the downregulated targets, we identified genes downstream of BCR, genes downregulated after PI3K inhibitor treatment, cell cycle, mitosis, DNA repair, and aurora kinase A.

The downregulation of BCR-related genes and upregulation of GPCRs was confirmed by an upregulation at the protein level of p-AKT and a downregulation of p-BTK (Supplementary Figure 4).

To look for differences or similarities of pathways modulated between Cluster A and Cluster B, we performed an enrichment map analysis using Cytoscape software (Supplementary Figure 5). Despite the differences in gene expression modulation, the two clusters converged toward using similar pathway changes. This could suggest two convergent mechanisms of resistance.

We identified pathways associated with secreted factors among the gene sets upregulated in both clusters. We have previously documented that secreted factors such as IL-6 and IL-16 are involved in lymphoma drug resistance^31,32^; Therefore, we treated parental cell lines with TK216 in the presence of conditioned media from resistant cell lines, but no effect was observed (Supplementary Figure 6).

Taking advantage of the RNA-seq data, we also analyzed parental and resistant cells’ mutational status and confirmed the cluster groupings observed by gene expression (Figure S7A). Focusing only on missense mutations with protein-coding implications, we identified ESR2, USP24, LRCH1, FAM200A, SFSWAP, and TRAPPC11 commonly mutated in resistant clones 2 and 5; FCGR2B, SRSF11, and PATJ were mutated in resistant clones 3 and 4 (Supplementary Figure 7B).

### Pharmacological screening of resistant clones

To identify compounds with different sensitivity between resistant and parental cells, we performed a pharmacological screening with a library of 348 compounds in resistant clone 2, as representative of Cluster A, and resistant clone 3, as representative of Cluster B, compared to one parental cell line. As previously reported, the library included approved kinase inhibitors, epigenetic compounds, and small molecules targeting critical biological pathways, such as PI3K/AKT/MAPK signaling and apoptosis (Supplementary Table 2) ^33^. Cells were treated with two concentrations of each library compound (500 nM and 50 nM) for 72 hours. Compounds associated with 15% or higher difference in cell proliferation between parental and resistant cell lines were selected. Despite belonging to two different clusters, the two resistant clones behaved similarly, in agreement with the similarities observed in deregulated pathways.

Compounds with more potent activity in resistant clone 2 compared to parental cells included inhibitors of BCL2, PI3K/AKT, topoisomerase, PIM, and heat shock protein (HSP). Among the compounds showing reduced activity in the resistant clone, there were inhibitors of aurora kinase, HIF, HDAC, VEGFR, multi-kinase inhibitors, and 2-Methoxyestradiol (Figure 4A-B, Supplementary Table 2).

**Figure 4.**
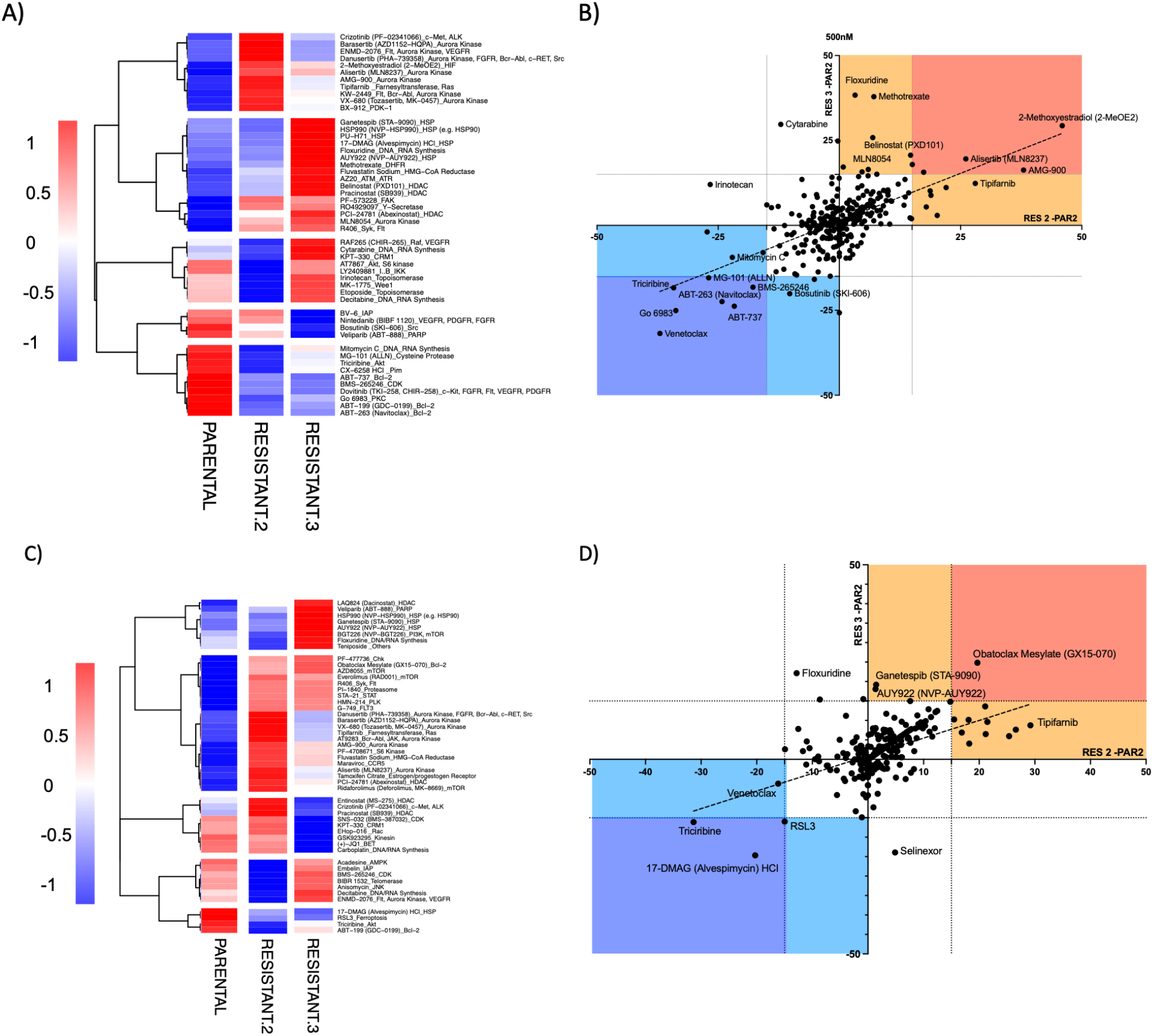
Pharmacological screening in TK216-resistant clones. Pharmacological screening was performed with library compounds 500nM (A, B) or 50nM (C, D). In the heatmaps A) and C), the top 50 drugs with higher standard deviations are shown.

Clone 3 showed increased sensitivity to inhibitors of BCL2, AKT, VEGFR, IAP, HSP, and XPO1. On the contrary, among the compounds showing less activity in the resistant clone, we identified inhibitors of Aurora Kinase or multi-kinase inhibitors, DNA/RNA synthesis, HSP, HDAC, and ATM/ATR (Figure 4C-D, Supplementary Table 2).

Finally, we identified anti-metabolites (antifolates and pyrimidine analogs) and compounds targeting aurora kinase as less active in resistant cells, in agreement with the downregulation of mitosis and cell cycle genes and aurora kinase A-related pathways (Supplementary Table 1).

The observation that BCL2-targeting compounds were more active in resistant cells than in parental ones was consistent with prior evidence showing venetoclax as one of the few compounds synergizing with TK216 in DLBCL cells ^13^. We confirmed the higher activity of BCL2 inhibition with venetoclax in resistant clone 2 cells, via a dose-response experiment (Figure 5A). The increased sensitivity in resistant clone 2 reflected a baseline status of antiapoptotic proteins like venetoclax-treated cells ^13^, with decreased BCL2 and increased MCL1 (Supplementary Figure 8). Surprisingly, the pan-BCL2 inhibitor obatoclax was less active in both resistant cells. Since MCL1 is one of the different targets between obatoclax and the other molecules that had increased activity in the resistant cells, we assessed the activity of the MCL1 inhibitor S63845 as a single agent in resistant and parental cells and in combination with TK216 (Figure 5A). However, the MCL1 inhibitor S63845 showed increased activity in both resistant clones (2 and 3), which did not explain the opposite activity observed between BCL2 inhibition and obatoclax. In addition to that, the combination of TK216 and S63845 showed an overall benefit in parental and resistant cells (Figure 5B-C).

**Figure 5.**
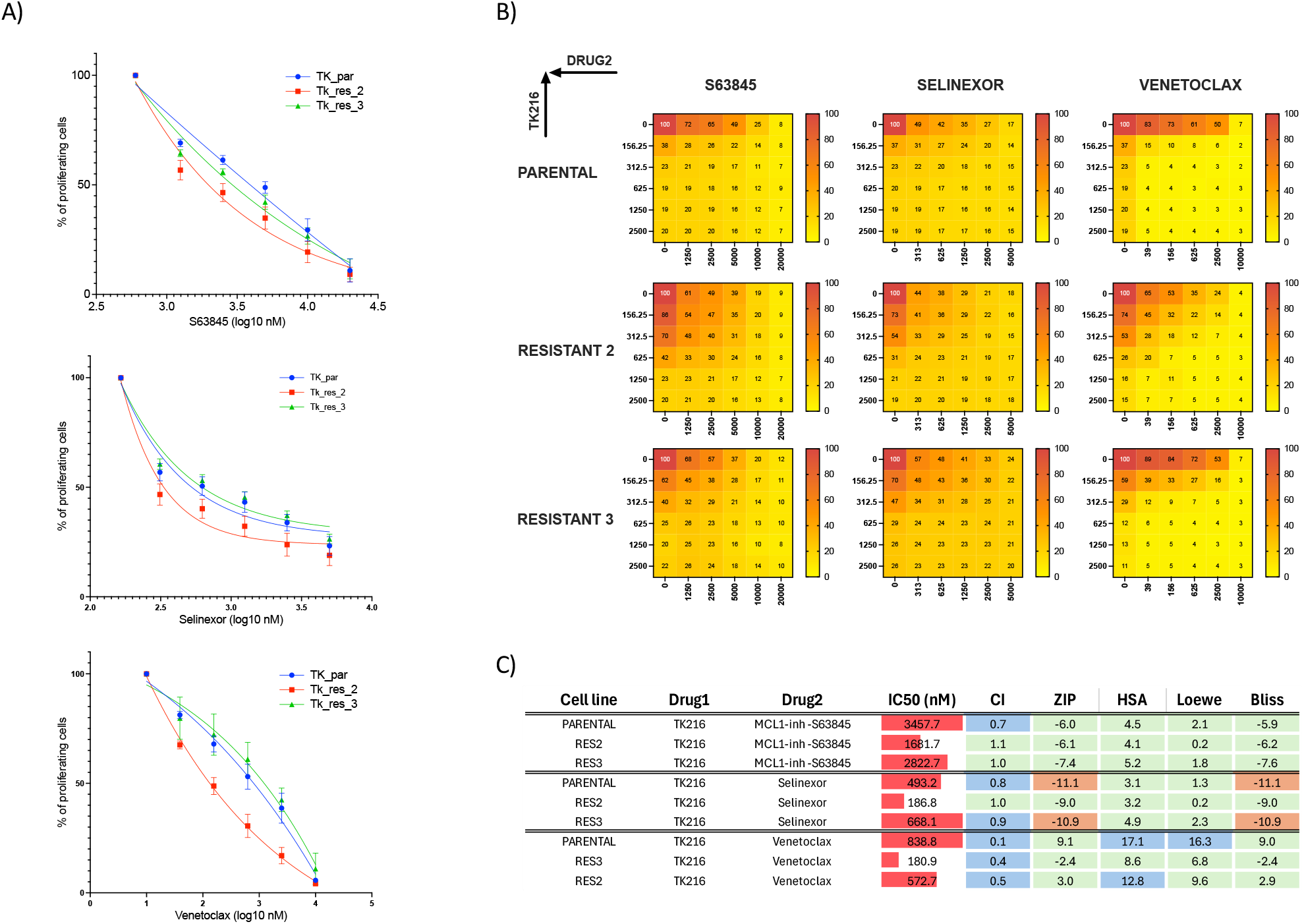
Single and combined treatment of venetoclax, S63845, and Selinexor in resistant clones 2 and 3 compared to parental 1. A) Dose-response curve of venetoclax, S63845, and selinexor in resistant 2/3 and parental cell lines. B) Heatmaps showing the % of proliferating cells in combination experiments. C) Table summarizing IC50s and combination values. Synergy: CI < 0.9; Additivity: 0.9 < CI <1.1; No benefit/antagonism: CI > 1.1. Synergy: ZIP/HAS/Loewe/Bliss >10; Additivity: -10 < ZIP/HAS/Loewe/Bliss <10; No benefit/antagonism: ZIP/HAS/Loewe/Bliss < - 10.

From the screening, the CRM1/XPO1 inhibitor selinexor showed higher activity in resistant clone 2 but not in 3, which was confirmed in a dose-response experiment (Figure 5A). In our previous work, this compound, FDA-approved for relapsed/refractory DLBCL patients ^34^, had an additive effect in two of four DLBCL cell lines tested, which did not include U2932. Here, we tested selinexor as a single agent in resistant and parental cells and in combination with TK216 in the U2932 (Figure 5). Even though we observed an increased sensitivity of selinexor in resistant clone 2 compared to parental cells as a single treatment, we did not observe such a difference in combination with TK216. Selinexor showed a benefit or synergism when combined with TK216 in parental and resistant clones.

## DISCUSSION

ETS-targeting agents have shown preclinical antitumor activity in various ETS-driven tumors, including DLBCL ^13,35,36^, and TK216 is a first-generation ETS-targeting small molecule that completed a Phase 1 trial for Ewing sarcoma patients. To identify resistance mechanisms that could be faced in patients, we developed five resistant clones through prolonged treatment with TK216, and we explored established anti-cancer drugs that mitigate the resistance to TK216. One cell line, U2932, showed a range of changes that could be responsible for resistance. The magnitude of resistance, from 150 nM (sensitive) to 1 µM (resistant), is important since drug levels in the Phase 1 clinical trial achieved serum concentrations above that with limited toxicity^15^. However, a second cell line, DOHH2, did not establish stable resistance to TK216.

Starting from an ABC-DLBCL cell line U2932, which is TK216 sensitive ^13^, we obtained resistance in five out of 24 wells of cells exposed to the IC90 of the ETS-targeting small molecule for almost two months. The cells were then kept for six months under TK216, and the resistance was stable after two weeks of washout, which was caused by three different mechanisms. One modality, represented by the resistant clone 9, was the multidrug resistance pump efflux protein MDR1 (ABCB1) overexpression. The overexpression was seen in RNA-seq data and confirmed by real-time PCR and immunoblotting. MDR1 is an ATP-dependent efflux pump that drives resistance to many drugs through an increased efflux of drugs out of the cells. Resistant clone 9 showed increased resistance to TK216, YK-4-279, and vincristine, a well-known MDR1 substrate ^29^. These data indicate that TK216 and YK-4-279 are substrates of MDR1, whose overexpression can be a resistance mechanism. In the remaining four clones, only two (2 and 3) showed a modest increase in MDR1 expression but did not show a substantial difference in vincristine sensitivity. This demonstrates that MDR1 did not mediate their resistance to TK216 and did not represent the only way cells could escape from TK216.

The other resistant clones were divided into two clusters based on their transcriptome profiling and mutational landscape: A (clones 2 and 5) and B (clones 3 and 4). The two clusters had different upregulated and downregulated genes. However, these converged to similar pathways, likely relevant to surviving the exposure to TK216. In all these clones, we saw an upregulation of genes upregulated by BET inhibitors, genes modulated by SPIB and IRF4, and a downregulation of genes related to DNA repair, cell cycle, and splicing. Among the genes mutated in cluster A, we identified ESR2 (estrogen receptor), ubiquitin-specific protease USP24, the negative regulator of CDC42 (LRCH1), and the alternative splicing factor (SFSWAP). In contrast, in cluster B, we identified the splicing factor SRSF11 and PTAJ involved in localizing proteins to the cell membrane. These mutations must be validated for their potential contribution to resistance to TK216. This is particularly the case for genes such as SRSF11 and SFSWAP, considering the role of alternative splicing and the participation of some ETS factors in splicing mechanisms ^37^.

Performing a high-throughput drug screening with a library of 348 FDA-approved compounds, we have identified compounds whose activity changed in resistant cells compared to the parental ones. Among the compounds more active in resistant cell lines, we identified BCL2 inhibitors, which were true for resistant clone 2 as a representative of cluster A. The difference in BCL2 inhibitor sensitivity correlated with different baseline expressions of BCL2 and MCL1. This was in perfect agreement with the fact that the BCL2 inhibitor venetoclax was one of the few compounds synergizing with TK216 in our previous work ^13^. Resistant clone 2 also showed an increased sensitivity compared to parental cells to CRM1/XPO1 inhibitor selinexor and MCL1 inhibitor S63845. Resistant clone 3, representative of cluster B, showed increased sensitivity to the MCL1 inhibitor S63845. Despite the increased sensitivity to these compounds, we could not appreciate an increased synergism when those compounds were combined with TK216. Of interest, despite increased sensitivity to venetoclax (BCL2), navitoclax (BCL2 and BCL-XL), and S63845 (MCL1), we observed reduced activity of obatoclax (BCL2, BCL-XL, and MCL1), suggesting that the resistant cells likely underwent a reprogramming of apoptotic dependencies, becoming reliant on specific anti-apoptotic proteins like BCL2 or MCL1 rather than a broad set of these proteins. This reprogramming might reduce their sensitivity to broad inhibitors like obatoclax but increases vulnerability to more selective inhibitors targeting the proteins on which they now depend for survival.

Recent research has shown that YK-4-279 synergized with Ewing sarcoma cells and the chemotherapeutic drug vincristine^38^. Prior work showed YK-4-279 resistance with some changes in gene expression; however, these specific genes were not evaluated as resistance mechanisms {Conn, 2020 #295}. Developing resistance to TK216 in an Ewing Sarcoma cell line previously engineered to make it DNA mismatch repair deficient, Povedano and colleagues identified microtubules as potential targets of TK216 ^39^; importantly, their experiments did not evaluate the drug’s effect on EWS-FLI1. We did not identify mutations in the gene coding for α-tubulin, as they reported ^39^. However, one of the compounds with the most reduced activity in our resistant DLBCL cells was 2-Methoxyestradiol (2-ME2), which can inhibit tubulin polymerization by interacting at the colchicine site ^40^, indicating that tubulin deregulation might still contribute to the resistance to TK216.

In conclusion, we have shown that multiple mechanisms might sustain the development of resistance to TK216 in DLBCL cells exposed to high doses of the compound. Possible mechanisms include the overexpression of the multidrug resistance pump efflux protein MDR1 and activation of multiple anti-apoptotic proteins. However, none of the 24 DOHH2 replicates developed stable resistance to TK216, suggesting that resistance may be cell line or lymphoma subtype-specific. Fortunately, resistance levels were below the achieved concentrations of TK216 in patients. Future considerations for clinical trials of TK216 include adding BCL2 or CRM1/XPO1 inhibitors, and their implementation in the next generation of clinical trials might overcome possible resistance.

## Acknowledgments

This work was supported by funding from the Leukemia & Lymphoma Society #6521-17 (to F. Bertoni and J.A. Toretsky). J.A. Toretsky holds the Hyundai Hope on Wheels Professorship in Pediatric Oncology.

