## Supplementary tables and figures for "Exploring Resistance to ETS Targeting Agents in Diffuse Large B-Cell Lymphoma"

**Supplementary Table 1. Transcriptomic signature distinguishing TK216-resistant cluster A and B versus parental cells.** The list shows log2 fold change (resistant vs parental) and P values for all quantified genes. Negative values indicate downregulation in resistance compared to parental cells, and positive values indicate upregulation. A two-tailed unpaired T-test was used.

**Supplementary Table 2. Compound sensitivity profile in TK216-resistant versus parental cells across a targeted inhibitor library.** The table reports percent viability relative to the vehicle for parental and two resistant cell lines representative of cluster A and B (resistant clone 2 for cluster A and resistant clone 3 for cluster B) at two concentrations (50 nM and 500 nM).

**Supplementary Figure 1. Dose-response curve of parental and TK216-resistant U2932 cells treated with Vorinostat.**

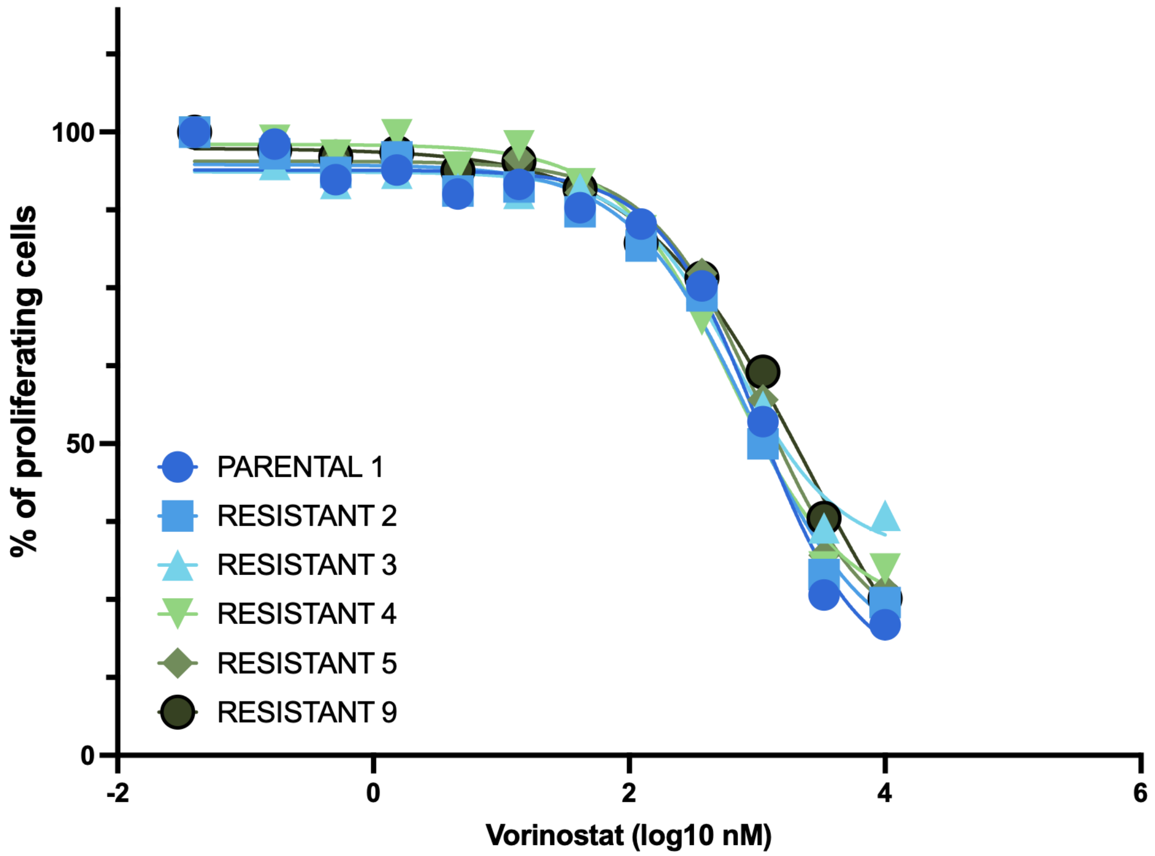

**Supplementary Figure 2. Vulcano plot of deregulated genes in Cluster A and Cluster B compared to parental cells.** A) Vulcano plot of deregulated genes in Cluster A compared to parental cells. B) Vulcano plot of deregulated genes in Cluster B compared to parental cells. The top 50 downregulated genes are highlighted in blue, and the top 50 upregulated genes are highlighted in red. Genes ranked based on FC with a minimal -log10 p value of 1.3

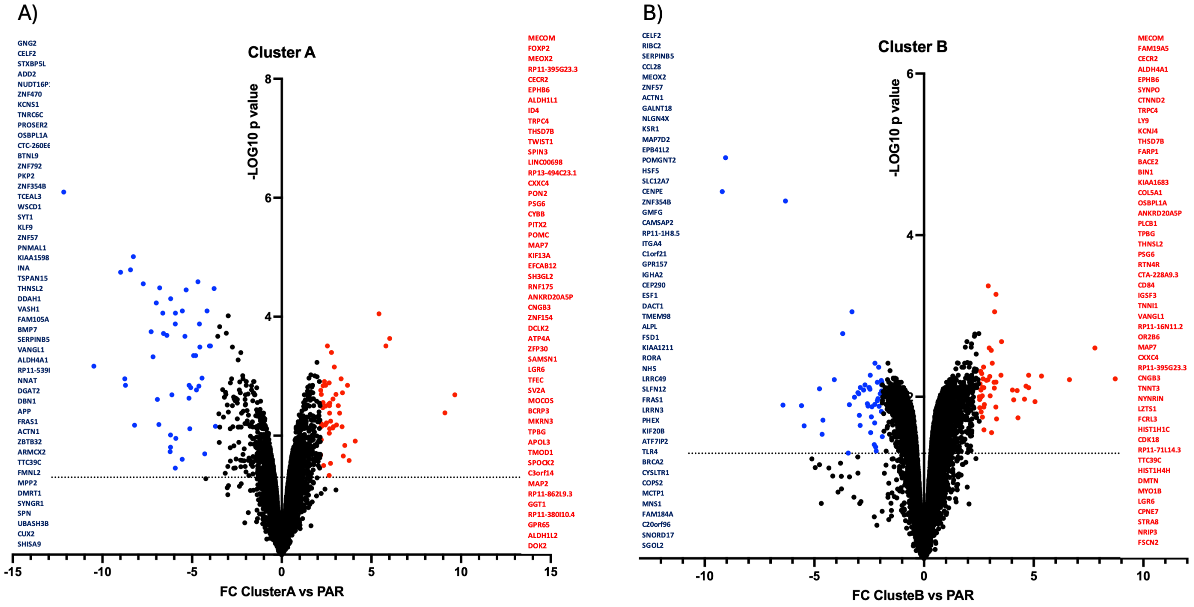

**Supplementary Figure 3. Gene sets upregulated or downregulated in resistant Cluster A and B (left and right panel, respectively).** GSEA, NES=normalized enrichment score obtained with gene-set enrichment analysis. Red bars = positive NES; Blue bars = negative NES. Upregulated gene-sets in the resistant Cluster A have positive NES, while downregulated genes have negative NES.

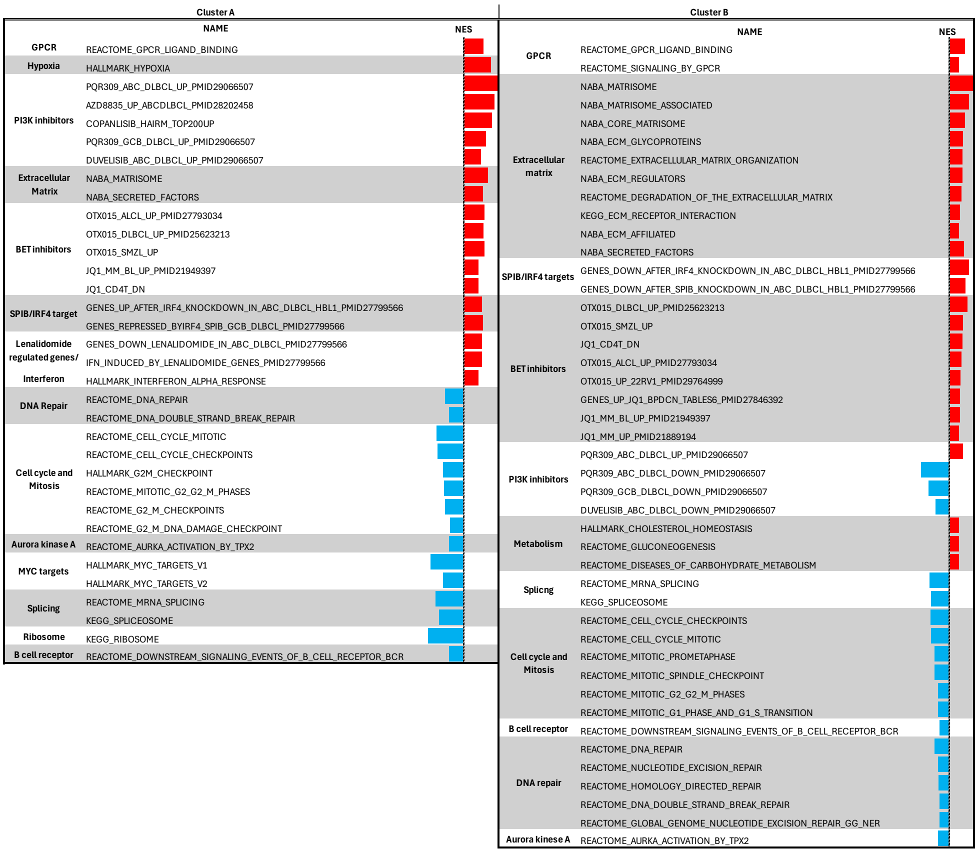

**Supplementary Figure 4. Immunoblot for resistant and parental cells with respective quantification.** Representative western blot of at least two biological replicates. M1 = membrane 1, M2 = membrane 2.

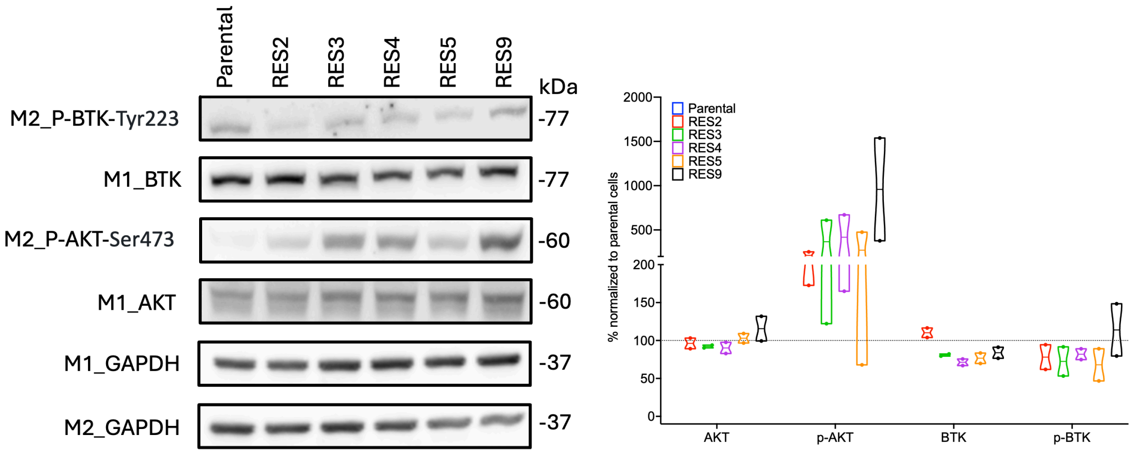

**Supplementary Figure 5. An enrichment map analysis comparing Cluster A and B resistant clones was performed.** An enrichment map analysis was performed with Cystoscape software to compare gene-set modulation between Cluster A and Cluster B. Every circle represents a gene set. Upregulated (red) or downregulated (blue) gene-sets in resistant compared to parental cells are shown in the left and right halves of the circle for Cluster B or Cluster A, respectively.

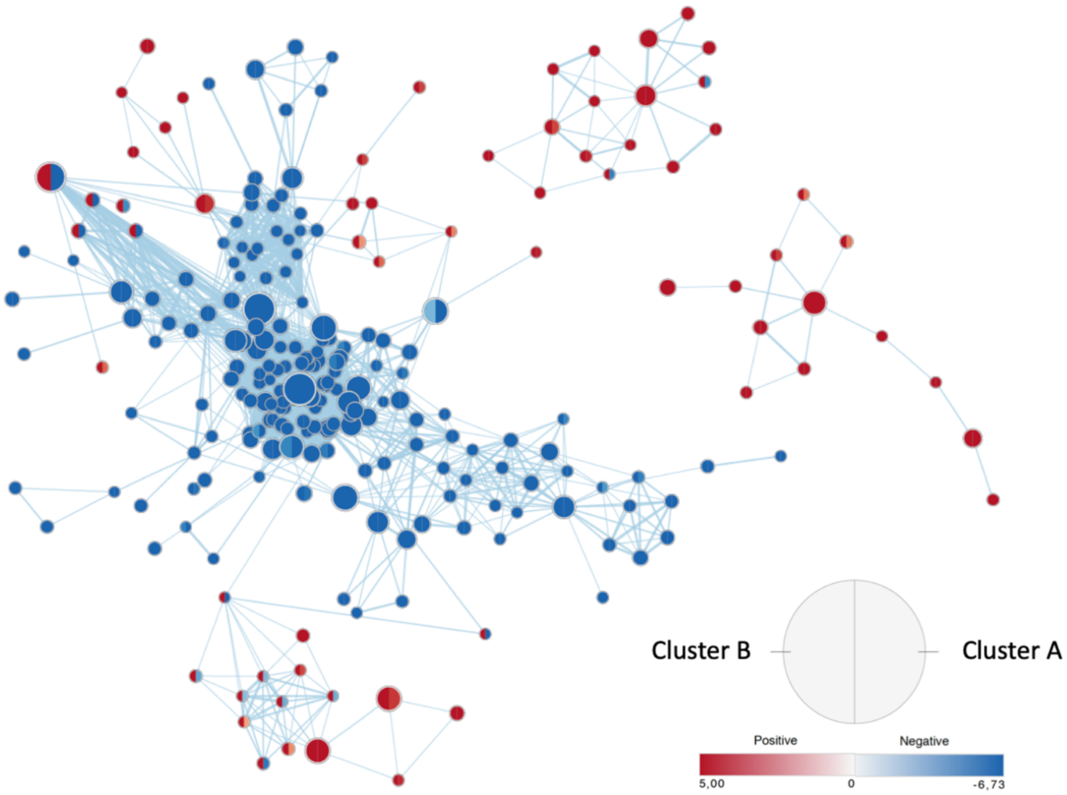

**Supplementary Figure 6. Dose response curve of parental cell line treated with TK216 in the presence of resistant cells' conditioned medium.**

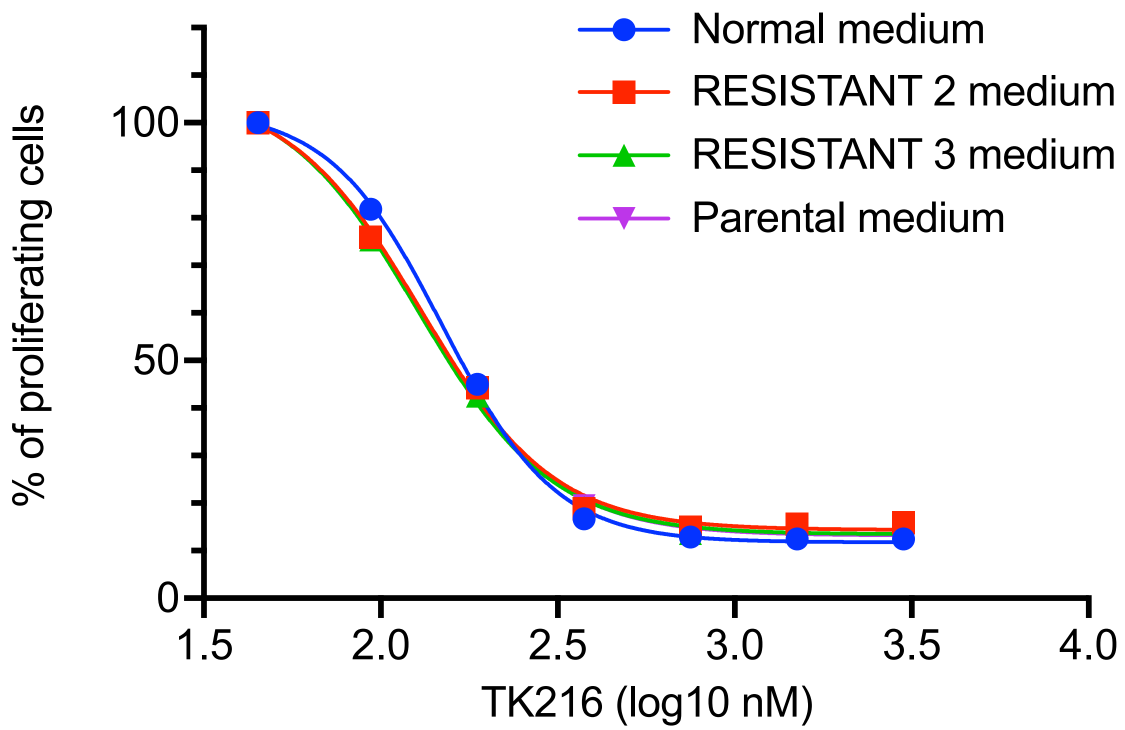

**Supplementary Figure 7. Resistant clones' mutational status.** A) Heatmap showing the mutation frequency in all genes differentially mutated between parental and resistant clones. Pearson's clustering method was used. B) The heat map shows genes with missense mutations that happen only in exons.

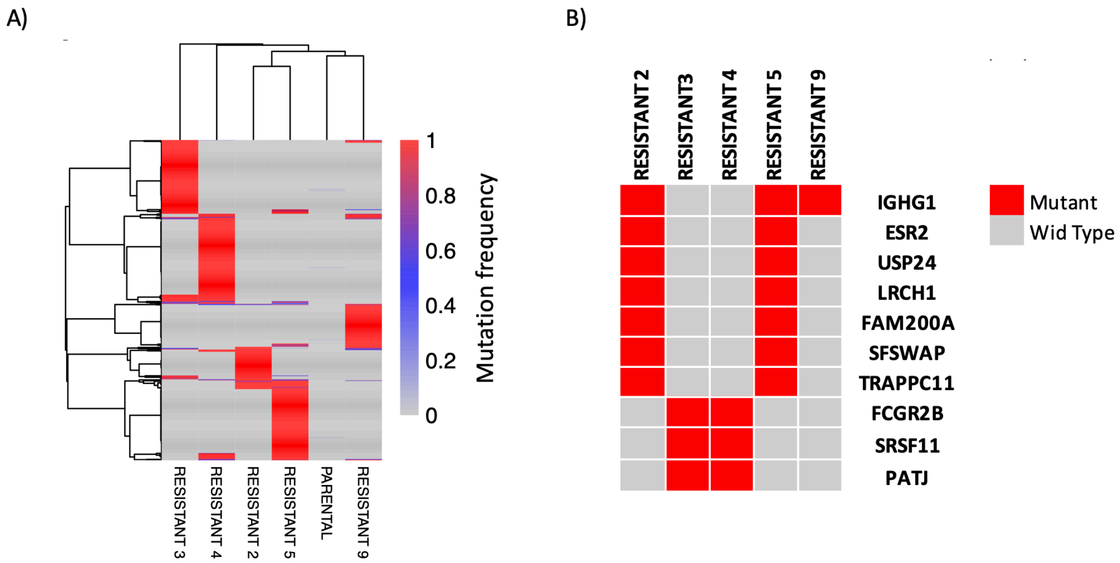

**Supplementary Figure 8. Immunoblotting of baseline expression of antiapoptotic proteins in TK216 parental and resistant clones.** Representative immunoblot of BCL2, Bcl-xL, and MCL1 protein expression. Vinculin was used as a loading control. M1 = Membrane 1; M2 = Membrane 2

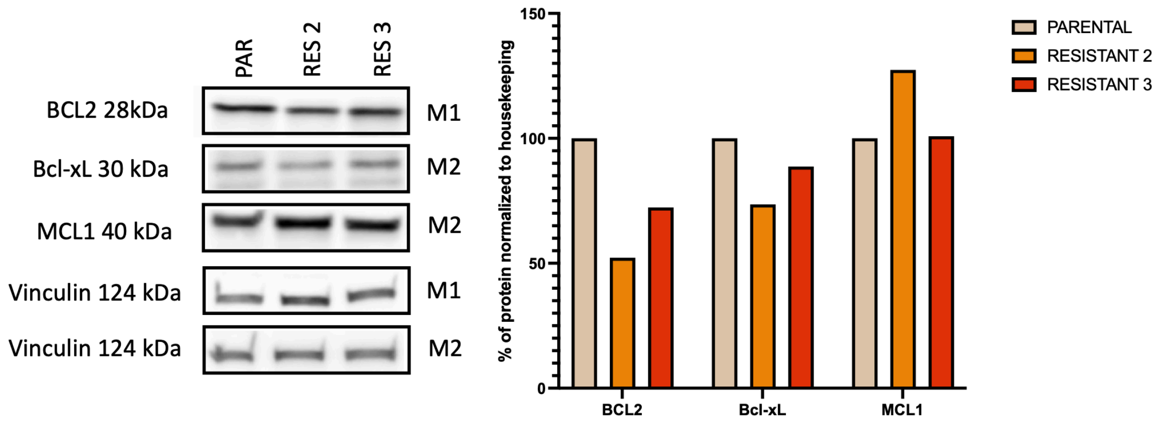
